# Ropinirole hydrochloride mitigates oxidative stress and neuroinflammation via the PI3K–mTOR pathway in TDP-43 hiPSC-derived microglial-like cells

**DOI:** 10.64898/2026.03.07.710283

**Authors:** Kagistia Hana Utami, Tatsuya Kozaki, Satoru Morimoto, Hirotaka Watanabe, Kensuke Okada, Nevin Tham, Shinichi Takahashi, Yasue Mitsukura, Hideyuki Okano

## Abstract

**Background:** Amyotrophic lateral sclerosis (ALS) is a fatal neurodegenerative disorder characterized by progressive motor neuron loss accompanied by neuroinflammation, oxidative stress, and impaired proteostasis, with pathological aggregation of TDP-43 as its defining hallmark. Microglia are recognized as central contributors to ALS pathogenesis, yet the mechanisms underlying their intrinsic dysfunction in the context of TDP-43 remain insufficiently characterized.

**Methods:** We characterized microglia-like cells (iMGLs) differentiated from isogenic TDP-43 M337V human induced pluripotent stem cell lines. We performed integrated functional assays, transcriptomic profiling and drug repurposing analysis to systematically compare mutant and control iMGLs. To assess therapeutic potential, we evaluated the effects of ROPI, a dopamine D2 receptor agonist previously advanced to clinical trials for ALS, on disease-relevant phenotypes in TDP-43 iMGLs.

**Results:** *TDP-43*^M337V/M337V^ iMGLs presented ALS-associated abnormalities, including cytoplasmic TDP-43 accumulation with impaired phagocytosis and elevated oxidative stress, impaired autophagy and mitophagy, altered cytokine profiles, and reduced ferritin levels. Building on our previous identification of ropinirole hydrochloride as neuroprotective for ALS motor neurons, we now show that it confers therapeutic benefits in mitigating microglial pathology. ROPI treatment significantly reduced oxidative stress and caspase-3/7 activity and partially restored cytokine homeostasis in *TDP-43*^M337V/M337V^ iMGLs, independent of autophagy/mitophagy modulation. Transcriptomic profiling revealed that ropinirole modulated disease-associated gene expression signatures involving protein folding, extracellular matrix organization, and oxidative stress responses. Furthermore, connectivity map–based analysis prioritized PI3K-Akt-mTOR inhibitors as candidates for reversing TDP-43 iMGL signatures, and ropinirole was found to modulate mTOR signaling. These converging lines of evidence support a mechanistic role for ropinirole in restoring microglial homeostasis via PI3K-mTOR pathway regulation. Taken together, our findings position ropinirole as a promising candidate dual action therapeutic candidate capable of targeting both neuronal and microglial dysfunction in ALS, suggesting broader applicability of ropinirole in modulating neuroinflammatory cascades across other neurodegenerative conditions.

**Conclusion:** Ropinirole broadens ALS therapy from motor neurons to microglia, underscoring its promise for integrated and clinically meaningful treatment strategies.

## Background

Amyotrophic lateral sclerosis (ALS) is a fatal neurodegenerative disease characterized by loss of motor neurons in the cortex, brainstem, and spinal cord, leading to paralysis and death, with a mean life expectancy of around four years after diagnosis [1]. ALS exhibit profound genetic and clinical heterogeneity, with more than 40 genes, including *SOD1*, *C9orf72*, *VCP*, *UBQLN2*, *FUS*, *TARDBP* (*TDP-43*), and *CHCHD10*, linked to both familial and sporadic forms [2–4]. Despite this complexity, nearly all cases converge on a common pathological endpoint: TDP-43 mislocalization and aggregation, observed in ∼97% of cases [2,5–8]. TDP-43 is a DNA/RNA-binding protein essential for RNA metabolism and gene regulation, ubiquitously expressed across all human tissues, but enriched in the central nervous systems, including neurons, oligodendrocytes and microglia [9–12].

Neuroinflammation is a pervasive feature of ALS, present in both sporadic and familial forms, and tightly linked to motor neuron degeneration. Across postmortem tissue, animal models and neuroimaging studies, ALS is marked by widespread activation of microglia and astrocytes, infiltration of peripheral immune cells and elevated proinflammatory cytokines, underscoring the critical role of non-neuronal immune responses in disease progression [13–18].

Microglia are tissue-resident macrophages of brain parenchyma with diverse roles in phagocytosis, synaptic pruning, and vascular regulation. In response to injury and neurodegeneration, they undergo context-dependent changes in gene expression, morphology and function, shifting from a protective to a reactive, neurotoxic state as ALS progresses [13,19,20]. This dual phenotype contributes to motor neuron loss and positions microglia as a compelling therapeutic target, though anti-inflammatory strategies have yielded only modest survival benefits [17,21,22]. Altered microglial phenotypes have been observed in postmortem ALS tissue and *Sod1* mouse models, where they modulate disease progression[23–27]. Studies using an inducible hTDP-43ΔNLS8 mouse model further demonstrated that TDP-43 expression increases the number of reactive microglia, exerting neuroprotective functions by selectively clearing neuronal TDP-43 aggregates. Conversely, when microgliosis is inhibited, motor recovery is impaired, suggesting critical roles of microglial dynamics in ALS[27,28]. While these studies implicate microglia in pathogenic processes during later stages of ALS, their contribution to early disease pathogenesis remains less well understood, with prior investigations largely relying on animal models, monocyte-derived macrophages, and postmortem human tissue, each limited by species-specific immune differences and limited temporal resolution[18,19,27–30].

Induced pluripotent stem cell (iPSC)-derived microglia-like cells (iMGLs) more faithfully recapitulate brain-resident microglia than monocyte-derived MGLs, preserving key transcriptional and functional signatures to study cellular pathophysiology in human-specific context. Recent work has revealed disease-specific phenotypes in C9orf72 [31–33], FUS [34], VCP [35], PFN1 [36] mutant iMGLs. One study using monocyte-derived microglia like cells from sporadic ALS patients showed striking neuroinflammatory profile [18], including cytoplasmic localization of TDP-43, impaired phagocytosis and altered cytokine profiles. However, investigations into iPSC-derived iMGLs within the context of TDP-43 mutational background remains unexplored. [18]

Here, we established isogenic hiPSC-derived microglia to model early neuroinflammatory processes in ALS using TDP-43 hiPSC-derived iMGL. Through functional assays, we uncover cell-autonomous changes in TDP-43 mutant microglia and demonstrate their relevance to ALS pathology. We previously identified Ropinirole (ROPI), a dopamine agonist originally developed for Parkinson’s disease, through iPSC-based motor neuron screens. ROPI has since been clinically trialed in ALS patients, demonstrating safety, tolerability, and delayed disease progression, thereby supporting its repurposing for ALS [14,15]. In this study, we show that ropinirole confers measurable benefits in attenuating microglial dysfunction, extending therapeutic actions of ROPI beyond motor neurons to include microglial pathology in ALS.

## Methods

### Cell lines

TDP-43 M337V heterozygous (*TDP-43* ^M337V/WT^) (JIPSC001108) and homozygous (*TDP-43* ^M337V/M337V^) (JIPSC001106) hiPSCs were generated via CRISPR/Cas9-mediated gene editing by Jackson Laboratories in the background of KOLF2.1J iPSCs (JIPSC001000) (https://www.jax.org/jax-mice-and-services/ipsc/cells-collection)[37].

### Maintenance of hiPSC cultures

Feeder-free isogenic hiPSCs were maintained in StemFit AKO2N media (Ajinomoto) under standard incubator conditions. The cells were dissociated into single cells via 0.5x TrypLE select (Cat no. 12563011) diluted in PBS and seeded at a density of 1.3 × 10^4^ cells/cm^2^ onto plates coated with 0.2 µg/cm^2^ iMatrix-511 (Laminin 511E8, Cat no. NP892-011; FujiFilm WAKO Pure Chemical Corp., Tokyo, Japan). The medium was changed every other day, and the cultures were passaged every 6–7 days.

### Differentiation of PSCs into iMGLs

To induce differentiation into iMGLs, we adapted the protocol previously developed by Sonn et al., with slight modifications[38]. Prior to initiating differentiation, the hiPSC colonies were dissociated via TrypLE select (0.5x in PBS), washed with PBS or DMEM/F12, and seeded at a density of 500 cells/cm^2^. Colonies were maintained in AKO2N media for approximately 5–7 days, after which they were induced for differentiation. The differentiation protocol consisted of hematopoietic progenitor cell induction via culture with freshly added cytokines during each medium change in medium 1 (StemPro34 Cat No. 10639011 (Thermo Fisher) supplemented with 500 µM L-ascorbic acid (Sigma, Cat. No. A4544), 200 µg/ml apo-transferrin (Nacalai-tesque, Cat No. 34401-55) and 450 µM monothiolglycerol (Sigma, Cat. No. M1753)). During the first five days of differentiation, the cells were incubated in hypoxic incubators. On Day 0, BMP4 (20 ng/ml, Peprotech Cat. No. 120-05) and CHIR99021 (3 µM) were added; on Day 2, BMP4 (20 ng/ml, Peprotech Cat. No. 120-05), FGF-2 (20 ng/ml, Peprotech Cat. No. 100-18B) and VEGF (50 ng/ml, Thermo Fisher, Cat No. PHC9391) were added; on Day 4, VEGF (15 ng/ml) and FGF-2 (5 ng/ml) were added; on Days 6-12, VEGF (10 ng/ml), IL-6 (10 ng/ml, Peprotech Cat. No. 200-06), IL-3 (30 ng/ml, Peprotech Cat No. 200-03), FGF-2 (5 ng/ml), SCF (50 ng/ml, Peprotech Cat. No. 300-07) and DKK1 (30 ng/ml, Peprotech) or IWR1e (2.5 µM) were added; and on Days 12-18, FGF-2 (10 ng/ml), IL-6 (10 ng/ml), IL-3 (30 ng/ml) and SCF (50 ng/ml) were added. On day 18, floating iPSC-derived hematopoietic progenitor cells (iHPCs) were assessed for CD11b/CD45 expression via flow cytometry. The progenitor cells were collected between days 18 and 25 and cultured in medium 2 (IMDM:F-12 medium at a ratio of 3:1, supplemented with 1x N2 (Thermo Fisher, Cat no. 17502001) and 1x B27 with vitamin A (Thermo Scientific, Cat no. A3582801) and 0.05% BSA (Sigma)), followed by 21 days of maturation in medium 2 supplemented with M-CSF (50 ng/ml, Peprotech Cat. No. 300-25), IL-34 (100 ng/ml, Peprotech Cat. No. 200-34), and TGF-β (50 ng/ml, Peprotech, Cat. No. 100-21).

### Cell treatment for various assays

Ropinirole treatment: the iMGLs were allowed to mature for 3 weeks, and 1 week prior to the end of differentiation, ropinirole was added for 1 week. RNA was then extracted and analyzed via qPCR and RNA-Seq.

For the autophagic flux assay, iMGLs were matured for 21 days and treated with 100 nM bafilomycin A1 (Sigma`‒`Aldrich, Cat. No. B1793) for 6 hours on day 21. Following treatment, the cells were harvested and subjected to either immunoblotting or immunofluorescence analysis. For translational inhibition, cycloheximide (100 µg/ml; Sigma‒Aldrich, Cat. No. 01810) was added to the culture medium for 6 hours, after which the cells were collected for RNA extraction and subsequent qRT‒PCR analysis.

### RT‒qPCR analysis

Total RNA was isolated from harvested cell pellets via an RNeasy kit (Qiagen) according to the manufacturer’s protocol. For cDNA synthesis, a minimum of 500 ng of RNA was reverse transcribed via a high-capacity reverse transcription kit (Applied Biosystems). Quantitative real-time PCR (qRT`‒`PCR) was performed in triplicate, and 10 ng of cDNA, individual gene-specific primer pairs and TB Green Premix Ex Taq II (Bio-Rad Laboratories) were used. Thermal cycling and fluorescence detection were conducted on a Viia7 Real-Time PCR System (Applied Biosystems, Thermo Fisher). Relative gene expressions were calculated via the Ct method, and GAPDH was used as the endogenous reference gene. A complete list of primers can be found in Table S1.

### Cytokine profiling

To assess cytokine secretion, 3-week-old mature iMGLs were seeded into 96-well plates at a density of 50,000 cells per well in 100 µl of medium. The cells were allowed to acclimatize overnight, followed by treatment with 100 ng/ml lipopolysaccharide (LPS) for 24 hours to induce inflammatory activation. After incubation, the cultured supernatants were collected and assayed via multiplex bead-based ELISA (LEGENDplex Human Inflammation Panel 13-plex; Biolegend Cat no. 740808) in accordance with the manufacturer’s instructions. Data acquisition and analysis were performed via CytoFlex (BD Biosystem) and the LEGENDplex Data Analysis Software suite.

### Incubation assay

To assess ROS generation, iMGLs were seeded onto 96-well black, clear bottom plates at a density of 500 cells/cm^2^. Following a 3-week iMGL maturation period under standard culture conditions, the cells were incubated with 5 μM CellROX DeepGreen Reagent (Thermo Fisher Scientific, Cat no. C10444) according to the manufacturer’s protocol for longitudinal live-cell imaging. The fluorescence was monitored via IncuCyte every 2-hour interval for 48–96 hours to quantify the ROS dynamics. For Caspase3/7 activity, the cells were incubated with CellEvent Caspase3/7 (Thermo Fisher Scientific, Cat no. C10430) Red diluted 1:100 in medium and incubated for 30 minutes prior to recording, according to the manufacturer’s protocol. Fluorescence was monitored via IncuCyte every 2-hour interval for 48 hours to monitor Caspase3/7 activity.

For the phagocytosis assay, iMGLs were seeded onto 96-well black wall–clear bottom plates at a concentration of 300–500 cells/cm^2^. The cells were allowed to mature for 3 weeks prior to the addition of 5 µg of pHRodo Red Zymosan Bioparticles (Thermo Fisher, Cat no. P35364). Phase contrast and fluorescence images were acquired every 30 min for 24 h via an IncuCyte S3 live-cell analysis system (Essen Bioscience). Analysis of fluorescence intensity was performed via the IncuCyte analysis software suite IncuCyte Zoom. Total intensity measurements were averaged from 4 separate fields from quadruplicate wells.

### Immunoblot

*The* iMGLs were pelleted via centrifugation, lysed with RIPA buffer supplemented with complete protease inhibitor (Roche) and phosphatase inhibitor (Roche) on ice for 15 minutes, and centrifuged at 12,000 rpm for 15 minutes. The soluble fractions in the supernatant were collected as protein lysates and quantified via a BCA assay (Thermo Fisher). Approximately 500 ng of protein was loaded on each capillary of a JESS digital western system (ProteinSimple) according to the manufacturer’s instructions. The antibodies used are listed in Table S1. The signals were detected with HRP-conjugated secondary antibodies and visualized via Compass for SW software (ProteinSimple).

### Immunofluorescence

iMGLs were seeded onto 8-chamber glass slides or 13 mm coverslips placed on 24-well plates at a density of 300–500 cells/cm^2^. After 3 weeks of maturation, the cells were fixed with 4% PFA for 20 minutes at room temperature. The cells were washed with PBS and blocked for 1 hour with blocking solution (1% bovine serum albumin, 5% fetal bovine serum, 0.3% Triton-X, in PBS). The samples were incubated with primary antibodies in the same blocking solution overnight at 4°C. The primary antibodies used are listed in Table S1. After 3 washes in 1x PBS, the cells were labeled with the fluorescently labeled secondary antibodies Alexa Fluor-488 or 555 (Thermo Fisher) for 1 hour at RT and subjected to Hoechst staining for nuclear labeling. The slides were then mounted with Fluoromount (Thermo Fisher). Images were taken via a Zeiss LSM700 or an Olympus FV4000 and were analyzed via ImageJ software.

### Image quantification analysis

For nucleocytoplasmic ratio (N:C) quantification, nuclear regions were defined from PU1 staining, and cytoplasmic regions were manually outlined from TDP-43 staining and calculated by subtracting nuclear integrated density and area from cytoplasmic measurements, and mean fluorescence intensities (mean gray value) for each individual cell. The nucleus-to-cytoplasm (N/C) ratio was calculated as the nuclear mean intensity divided by the cytoplasmic mean intensity. For mitophagy quantification, colocalization between TOM20-labeled mitochondria and LC3B was quantified using Mander’s overlap coefficients (M1, M2) calculated with the Coloc2 plugin in Fiji/ImageJ.

### Flow cytometry

Between day 18 and day 25, floating microglial progenitors were collected and assessed via flow cytometry prior to being plated for maturation. Approximately 1/^4^ of the total iMGL cultures were analyzed by flow cytometry for each genotype. After centrifugation, the cells were stained with cell-surface marker-conjugated antibodies against CD45-FITC (eBioscience), CD43-APC (eBioscience), CD64-FITC (eBioscience) and CD11b-APC (BD Pharmingen) for 45 minutes on ice. Following washes with FACS buffer (PBS with 2% BSA), the cells were resuspended in FACS buffer and analyzed via a Beckman Coulter CytoFlex machine.

### Bulk RNA sequencing

*The* iMGLs were allowed to mature for 3 weeks, after which RNA was extracted from 3 controls, 3 TDP-43^M337V/M337V^ cultures, with and without ropinirole treatment using the RNeasy Mini Kit (Qiagen) according to the manufacturer’s instructions. Ropinirole (10 µM) treatment was administered during the final 7 days of iMGL maturation, after which samples were collected and processed for bulk RNA-seq. Library preparation and paired-end 150 bp sequencing were subsequently performed, and 20 million reads/sample were obtained via NovaSeq system.

The RNA-seq data were analyzed via R and RStudio. Detailed quality control of aligned reads was assessed via FastQC [39]and MultiQC v.1.12[40]. Low quality reads were removed by setting a minimum length of 35 bp and a minimum Phred quality score of 25. RNA-seq reads were aligned to the human reference genome GRCh38.p12, and mapped with Salmon[41]. Gene expression quantification was performed with feature counts. The statistical analysis started by filtering genes with low expression levels via HTS Filter. Differential gene expressions were calculated with DESeq2 [42]at the gene level with R v.4.2.0. Results contrasts were generated by 3 layers of comparisons: untreated TDP-43 versus untreated isogenic control, Ropinirole treated TDP-43 vs untreated TDP-43 untreated, and Ropinorole treated TDP-43 versus Ropinirole treated Control. Genes were considered differentially expressed at LFC 1, FDR < 0.05. Significantly up and downregulated differentially expressed genes were used as input to functional overrepresentation analyses to identify enriched pathways using EnrichR[43] and clusterprofiler4.0[44]. ClusterProfiler searches the following data sources: Gene Ontology (GO; molecular functions, biological processes, and cellular components), KEGG, WikiPathways. ClusterProfiler reports the hypergeometric test p-value with an adjustment for multiple testing using Bonferroni correction. Overrepresented function categories are plotted in cnetplot, where the top significant terms were visualized in node-based network pathway.

### Data availability

Bulk RNA sequencing and processed data generated during this study were deposited at the Gene Expression Omnibus of the National Center for Biotechnology Information under accession number GSE288292 and will be available upon acceptance of the manuscript. All data and/or analyses generated during the study are available from the corresponding author upon reasonable request.

### Connectivity map analysis

We used the significant DEG results from the comparison of *TDP-43* ^M337V/M337V^ iMGL vs control iMGL to form the ALS-TDP-43 transcriptomic signatures to query the connectivity map (CMAP) integrated within the Library of Integrated Network-Based Cellular Signatures (LINCS) project from the National Institute of Health. For the ROPI signature, a transcriptomic comparison was performed between ROPI-treated iMGL and untreated TDP-43-iMGL cells. The gene log2fold changes from the differential expression analyses were further decreased via the normal DESeq2 estimator to derive the mutant and ROPI signatures. The ALS-TDP-43 signature was initially queried against drug signatures from THP-1 cells, a monocytic cell line derived from the peripheral blood of acute monocytic leukemia patients. THP-1 cells are widely utilized for studying macrophage function and are considered the cell line most closely related to the iMGL lineage. Signatures treated by drugs of unknown or unannotated MOAs were excluded from the query, and the Zhang scoring method (also known as statistically significant connections map ssCMap[45]) was used to calculate the connectivity score between a drug and the disease signature. Zhang scores can range from –1 to +1, where a score of –1 indicates an absolute reversal and a score of +1 indicates an exact phenocopy of the gene ranks and direction of regulation. In addition to the connectivity score, its empirical p value is also calculated to determine the significance of the score. An empirical p value smaller than 0.05 denotes a significant relationship between the two signatures (Table S4).

### Statistical analysis

Statistical analysis was performed via GraphPad Prism. The reported values are the means ± SEMs, noted in the figure legend for each panel. Unless otherwise noted in the Figure legends, n = 3 replicates for each genotype. Statistical significance was determined via Student’s t tests when two groups were compared or via ordinary one-way ANOVA with Tukey’s multiple comparison test when three or more groups were compared.

## Results

### TDP-43^M337V/M337V^ iPSCs efficiently differentiated into iMGLs

To capture early microglial function and activity in the context of the genetic background of TDP-43 mutation, we leveraged a CRISPR/Cas9-engineered hiPSC line carrying M337V variant in one (*TDP-43*^M337V/WT^) or both alleles (*TDP-43*^M337V/M337V^), located at the C-term domain of the TDP-43 protein (Fig. 1a). This well-characterized mutation, originally identified in familial ALS across three generations in U.S. and U.K. cohorts, is associated with slightly earlier disease onset and has been extensively characterized in mouse and iPSC-derived studies [46,47]. Parental wild-type KOLF2.1J cells were used as the isogenic control, as a widelycharacterized reference hiPSC line and has been used by NIH iPSC Neurodegenerative Disease Initiative [37,48–50]. All iPSC lines were differentiated into iMGLs using our previously published protocol with slight modifications [38] (Fig. 1b). The protocol involves the generation of hematopoietic progenitor cells and floating microglial precursors, which are then harvested, expanded and plated for maturation to reach a microglia-like state (Fig. 1c). We analyzed floating progenitors on day 18 of differentiation by flow cytometry, and the majority of these cells expressed CD45, CD11b, CD43 and CD235a (Fig. 1d). Maturing iMGLs were then assessed after three weeks of differentiation; these iMGLs expressed typical microglial-specific markers, such as *TMEM119* and *P2RY12* transcripts (Fig. 1e), as well as IBA1 (with a range of 67-83% IBA1+ cells and 80-90% P2RY12+ cells) (Fig. S1A). Notably, analysis of microglial marker expression identified significant reduction of *TMEM119* and *P2RY12* transcripts expression in *TDP-43*^M337V/M337V^ iMGL, suggesting an activated disease state in mutant iMGLs[51]. Thus, both engineered and control iMGLs were deemed suitable for investigating neuroinflammation in the context of TDP-43 mutation.

**Fig. 1.**
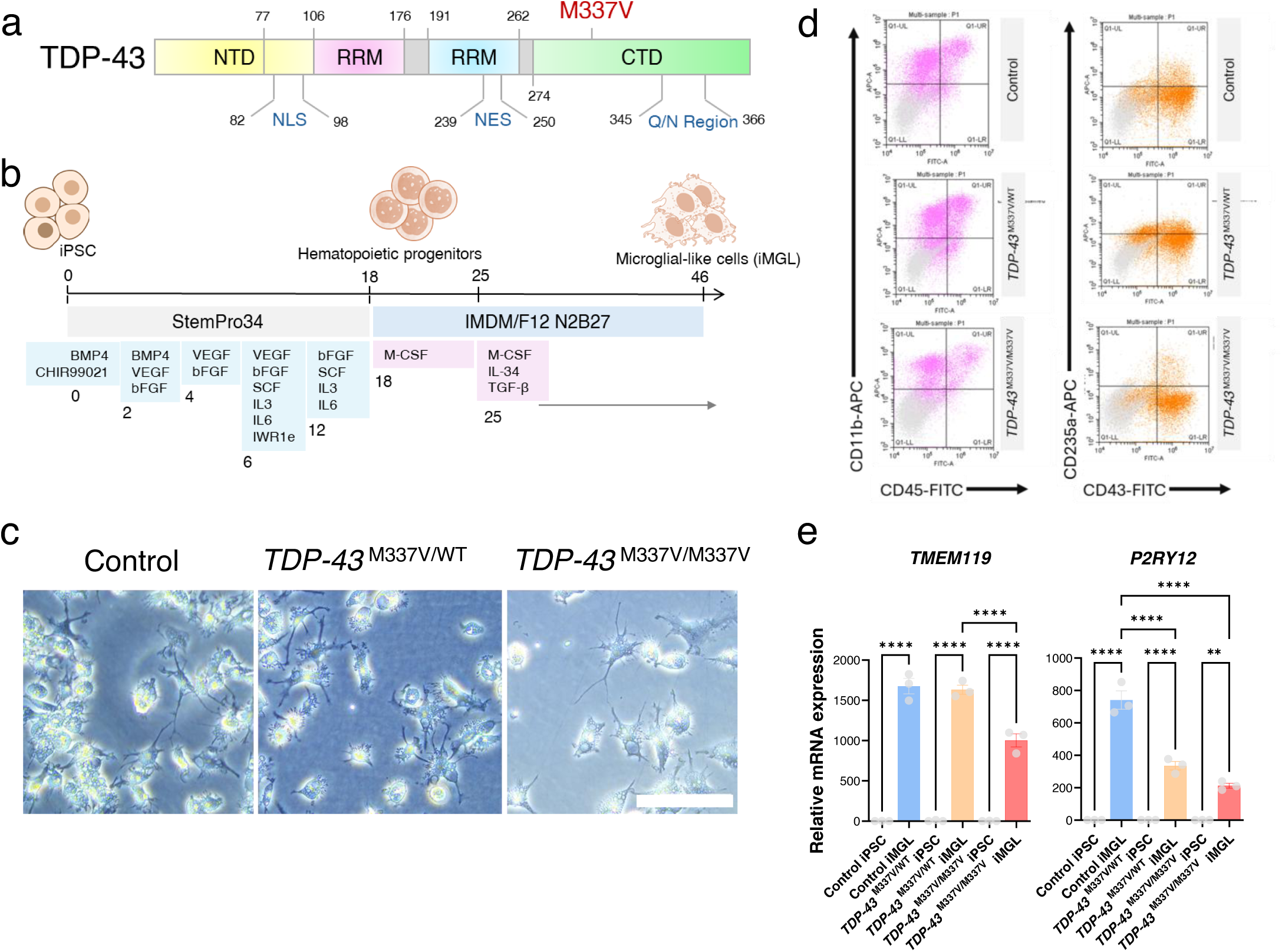
Generation and characterization of iMGLs from TDP-43 M337V-mutant and isogenic iPSCs. **(A)** Schematic representation of the TDP-43 protein domain. The M337V mutation is located at the C-term prion-like domain (PrLD). (**B**) Schematic representation of the protocol used to differentiate control and isogenic *TDP-43*^M337V/M337V^ iMGLs via a modified protocol, as described in the methods section. (**C**) Representative brightfield images of differentiated iMGLs upon maturation without any stimulation. Scale bar: 200 µm. (**D**) Flow cytometry analysis of cell-surface markers (CD11b, CD45, CD235a, and CD43) to characterize differentiated iMGLs on day 18 of differentiation. (**E**) iMGLs expressed the microglia-specific transcripts *P2RY12* and *TMEM119* upon maturation, as assessed by qRT`‒`PCR. The data represent three independent biological replicates for each genotype, and one-way ANOVA with Tukey’s multiple comparison test was used to determine significance (**** *P<0.0001, *** P* <0.001, ** *P <* 0.01, * *P <* 0.05).

Cytoplasmic mislocalization of TDP-43 is a hallmark of ALS, typically observed in post-mortem tissue of ALS patients. To assess whether this pathology extends to our iMGL model, we examined TDP-43 M337V iMGL and found cytoplasmic accumulation in both heterozygous and homozygous mutants with significantly increased nucleocytoplasmic ratio (Fig. 2a), in keeping to previously reported in the literature.

**Figure 2.**
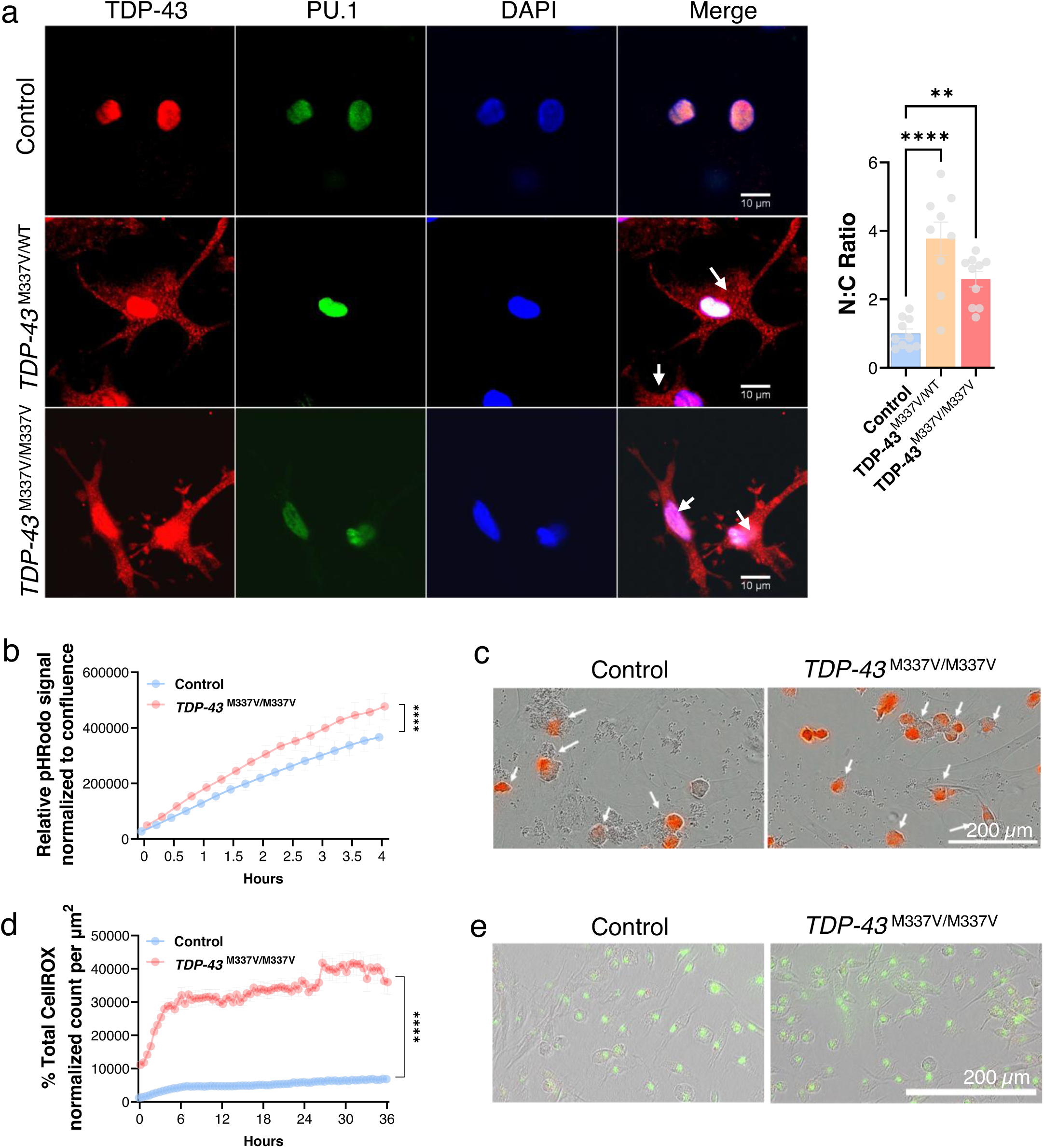
M337V iMGL shows altered TDP-43 localization and inflammatory activity. (A) Representative immunofluorescence images showing the subcellular localization of TDP-43 in the control and mutant TDP-43 groups, revealing the predominant nuclear localization of TDP-43 in the control group, whereas the mutant iMGLs presented the cytoplasmic localization of the TDP-43 protein. Scale bar: 10 µm. (Right panel): Quantification of nucleocytoplasmic (N:C) ratio from TDP-43 vs PU1 mean intensity values. (B) pHRodo zymosan uptake was normalized to the fluorescence intensity to measure phagocytosis. The data are presented as the means ± SEMs from quadruplicate wells, with four image fields per well. (**C**) Representative images of iMGLs at 6 hours after pH-induced rodent zymosan phagocytosis. The white arrows indicate engulfment of zymosan beads by phagosomes. Scale bar: 200 µm. (**D**) CellROX uptake was normalized to the fluorescence intensity to measure reactive oxygen species (ROS) production. The data are presented as the means ± SEMs from quadruplicate wells, with four image fields per well. (**E**) Representative images of iMGLs 6 hours after the CellROX labeling assay.

### Increased oxidative stress and reduced autophagic flux in TDP-43

We next investigated the functional consequences of TDP-43 mutation in microglial function. First, we determined that TDP-43 M337V mutant iMGL alters phagocytosis. Microglia constantly survey their microenvironment by continuously extending and retracting their processes in response to damage and disease, such as by phagocytosing dead cells. We determined that *TDP-43*^M337V/M337V^ iMGL under monoculture condition exhibited enhanced phagocytosis of zymosan A [52] (Fig. 2b, 2c, Fig. S1B), suggesting that *TDP-43*^M337V/M337V^ iMGLs possess an intrinsically primed or dysregulated phagocytic phenotype. This finding is in line with a recent immunophenotyping study on the human ALS postmortem brain, which revealed increased phagocytosis and the upregulation of genes indicative of increased phagocytosis[30,53]. Together, these data suggest that *TDP-43*^M337V/M337V^ iMGLs exert exaggerated phagocytic activity independent of neuronal debris, potentially reflecting a hyperreactive microglial state in ALS pathology as a consistent survival response.

Oxidative stress, driven by genetic mutations and additional environmental or cellular stressors, has been strongly implicated in ALS pathogenesis. However, the extent to which oxidative stress is altered in ALS patient-derived iMGL remains unclear. To assess the impact of TDP-43 mutation on the inflammatory state of microglia and oxidative stress levels, we assessed the production of reactive oxygen species (ROS) levels in iMGLs under steady-state conditions. *TDP-43*^M337V/M337V^ iMGLs had significantly increased ROS in absence of an inflammatory stimulus (Fig. 2d, 2e), indicating intrinsic susceptibility to oxidative stress as an inherent feature of ALS microglia and is not limited to motor neurons.

Autophagy is a crucial catabolic process that maintains cellular homeostasis by clearing protein aggregates and damaged organelles via lysosomal degradation. Activated under oxidative stress, it plays a central role in responding to ROS. We hypothesize that increased ROS levels in *TDP-43*^M337V/M337V^ iMGLs reflects impaired autophagic flux or mitophagy. To assess autophagic flux, we treated the cells with bafilomycin A1, a selective vacuolar (V)-ATPase inhibitor that blocks autophagosome`‒`lysosome fusion and lysosomal degradation, and measured LC3B, key marker for autophagosome formation. *TDP-43*^M337V/M337V^ iMGLs exhibited significant reduction of LC3B (Fig. 3a), suggesting impaired autophagic flux activity. We also evaluated mitochondrial clearance via mitophagy via immunofluorescence analysis of TOM20, a marker of the outer mitochondrial membrane, and LC3B. The colocalization of TOM20 and LC3B, as quantified by Mander’s colocalization coefficient, indicates the engulfment of mitochondria by autophagosomes, a key step in mitophagy. exhibited reduced mitophagy (Fig. 3b), further supporting the hypothesis that impaired autophagic and mitophagy processes contribute to oxidative stress in these cells.

**Fig. 3.**
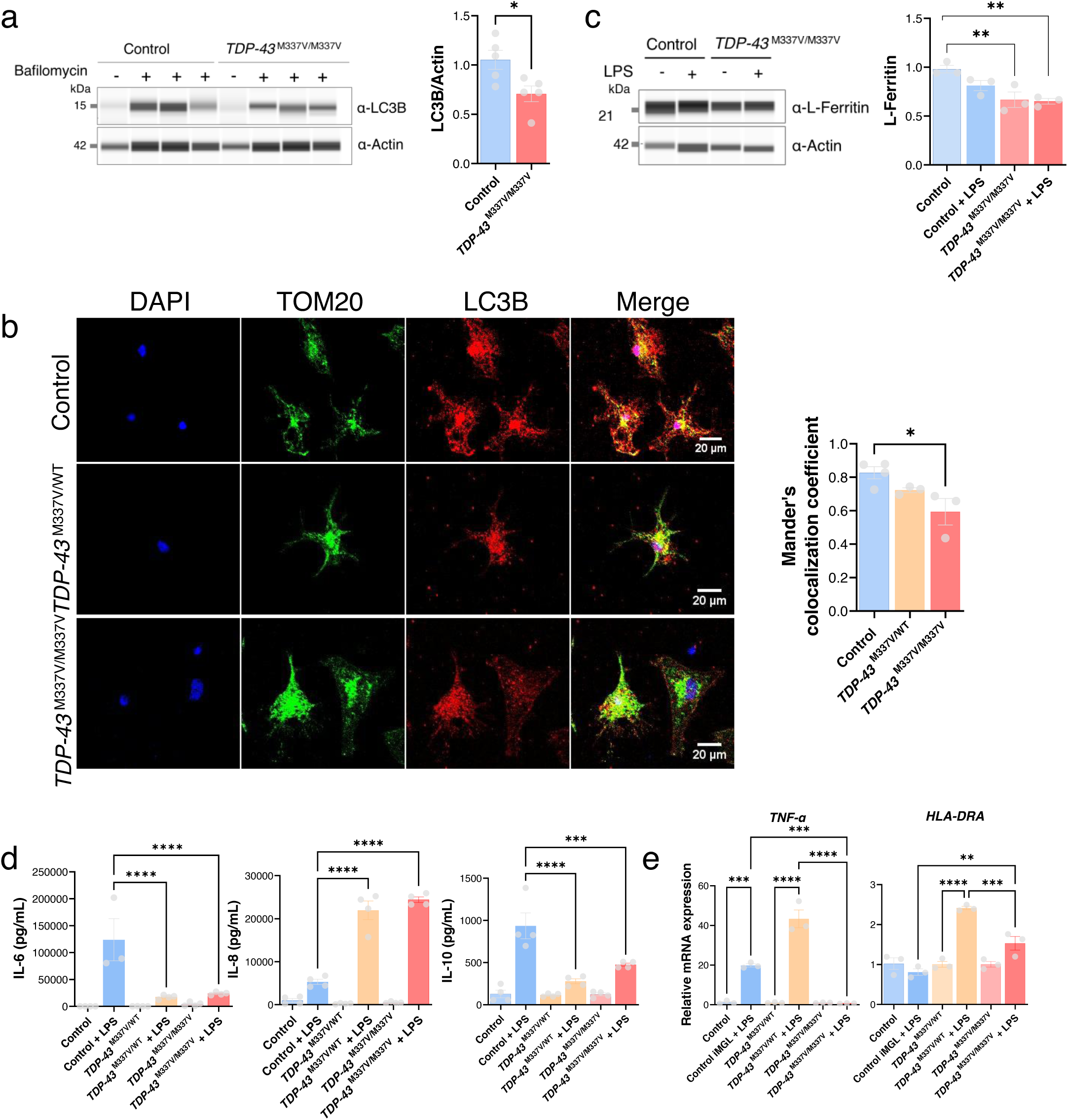
Characterization of the neuroinflammatory state of *TDP-43* M337V iMGLs. (**A**) Left: Untreated and bafilomycin-treated iMGLs were assessed by immunoblotting. Right: Quantification of bafilomycin-treated iMGLs expressing LC3B via immunoblot analysis. The presented values are the means ± SEMs (n = 3 biological replicates). (**B**) Left: Representative images of LC3B– and TOM20-expressing cells in mature iMGLs. Right: Quantification of LC3B and TOM20 colocalization coefficients from immunofluorescence images. The presented values are the means ± SEMs (n = 10 image fields from 2 coverslips per genotype). (**C**) Left: Representative immunoblot images of ferritin at homeostasis and upon microglial activation. Right: quantification of ferritin levels from immunoblot analysis. The data points are the means ± SEMs (n = 3 biological replicates per genotype). (**D**) Measurement of cytokine secretion with and without LPS stimulation via flow cytometry. The presented values are the means ± SEMs (n = 2 technical replicates from each 2 biological replicates from 2 differentiation batches). (**E**) qRT`‒`PCR analyses were performed to measure the transcript levels of cytokines that were not detected by the human inflammation panel. The data represent three independent replicates for each genotype from 2 different batches, and one-way ANOVA with Tukey’s multiple comparison test was used to determine significance.

Ferritin, an inflammation-regulated iron storage protein, is commonly elevated during disease progression and regulated by pro-inflammatory cytokines. We therefore measured L-Ferritin levels in iMGL and found significantly reduced L-Ferritin levels in *TDP-43*^M337V/M337V^ iMGLs (Fig. 3c), suggesting altered iron homeostasis and proinflammatory phenotype.

### Increased inflammatory responses in LPS-treated TDP-43^M337V/M337V^ iMGLs

Microglia respond to diverse stimuli with dynamic changes in morphology and gene expression. Upon activation, microglia can proliferate and release cytokines that exert either harmful or protective effects on neurons. To characterize inflammatory responses, we stimulated iMGLs with LPS and profiled cytokine secretion via flow cytometry and qRT-PCR. *TDP-43*^M337V/M337V^ iMGLs secreted lower levels of IL-6 and IL-10, and higher levels of of IL-8, *TNFα* and *HLA-DRA,* indicating a shift towards proinflammatory activity (Fig. 3d-e). Together, these results indicate that M337V variant in TDP-43 leads to dysregulation of the inflammatory responses in microglia.

### Ropinirole hydrochloride partially suppresses ALS-associated inflammatory phenotypes in TDP-43 iMGLs

Our group recently identified ROPI, a potential repurposed therapeutic for ALS, through a high-throughput phenotypic screen using iPSC-derived motor neurons [54,55]. Further studies have shown that ROPI has anti-ALS effects via D2R-dependent and D2R-independent mechanisms[55–57]. Given its antioxidant properties, we tested whether ROPI could alleviate inflammation and oxidative stress in *TDP-43*^M337V/M337V^ iMGLs. We treated iMGLs for one week and observed a marked reduction in ROS levels (Fig. 4a), indicating partial normalization of oxidative stress. To check for ROS-induced apoptosis, we measured Caspase3/7 levels and observed that elevated apoptosis in both *TDP-43*^M337V/M337V^ and control iMGLs were significantly reduced by ROPI treatment (Fig. 4b).

**Fig. 4.**
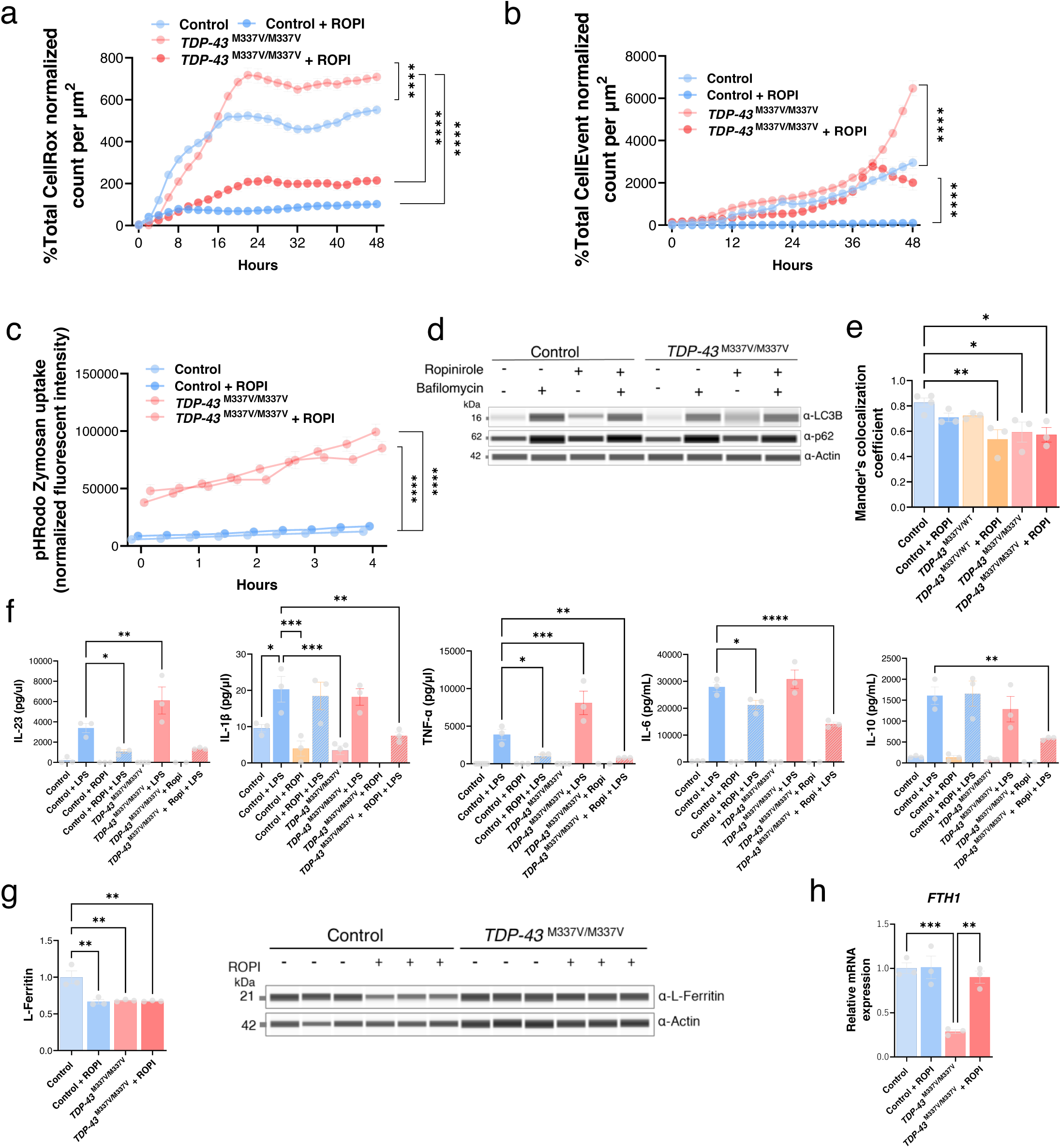
Ropinirole hydrochloride partially mitigates the neuroinflammatory phenotypes of TDP-43 iMGLs. (**A**) CellROX uptake was used to normalize the fluorescence intensity to measure reactive oxygen species (ROS) production in the presence of ROPI. The data are presented as the means ± SEMs from triplicate wells in four image fields. (**B**) CellEvent uptake was used to normalize the fluorescence intensity to measure Caspase3/7 activity in the presence of ROPI. The presented values are the means ± SEMs from triplicate wells and four image fields. (**C**) pHRodo zymosan uptake was normalized to the fluorescence intensity to measure phagocytosis in the presence of ROPI. The data are presented as the means ± SEMs from triplicate wells in four image fields. (**D**) Left: immunoblot analysis of autophagic flux using LC3B and p62 in the presence or absence of bafilomycin and ROPI. Right: quantification of autophagic flux in the presence of ROPI on the basis of immunoblot analysis. (**E**) Right: Quantification of LC3B and TOM20 Mander’s colocalization coefficients from immunofluorescence analysis for ROPI-treated and untreated iMGLs. (**F**) Measurement of cytokine secretion from ROPI-treated cells in the presence or absence of LPS via a human inflammation panel. The data points are presented as the means ± SEMs (n = 2 technical replicates from 2 biological replicates from 2 differentiation batches). (**G**) Immunoblot analysis and quantification of ferritin light chain expression, with or without ropinirole treatment. The presented values are the means ± SEMs from 3 replicates.

We next assessed ROPI’s impact on microglial function. ROPI treatment did not restore enhanced phagocytic activity in *TDP-43*^M337V/M337V^ iMGLs, as well as unable to normalize autophagy or mitophagy (Figure 4c-e). However, ROPI partially rescued cytokine secretion profiles, modulating neuroinflammatory responses (Fig. 4f). These protective effects on oxidative stress, apoptosis and cytokine output appear independent of autophagy and mitophagy. While ROPI did not normalize reduced L-ferritin protein levels (Fig. 4g), it restored *FTH1* transcript expression, indicating partial recovery of iron homeostasis (Fig. 4h).

### Molecular profiling of ROPI-treated ALS iMGLs revealed partial restoration of oxidative stress-related gene expression

To gain a deeper understanding of the global molecular changes induced by ROPI treatment, we performed bulk RNA sequencing on *TDP-43*^M337V/M337V^ iMGLs with and without ROPI treatment, together with their isogenic control pairs. The iMGLs were matured for three weeks, and ROPI was administered during week two to capture late-stage molecular changes while minimizing early differentiation effects.

Principal component analysis (PCA) revealed distinct clustering patterns based on genotype and treatment conditions (Fig. 5a). Interestingly, control iMGLs with and without ROPI treatments clustered closely within the PCA, whereas *TDP-43*^M337V/M337V^ clustered separately from the ROPI-treated *TDP-43*^M337V/M337V^. This suggests that ROPI induces a transcriptional shift in *TDP-43*-mutant iMGL with minimal effect on control iMGLs.

**Fig. 5.**
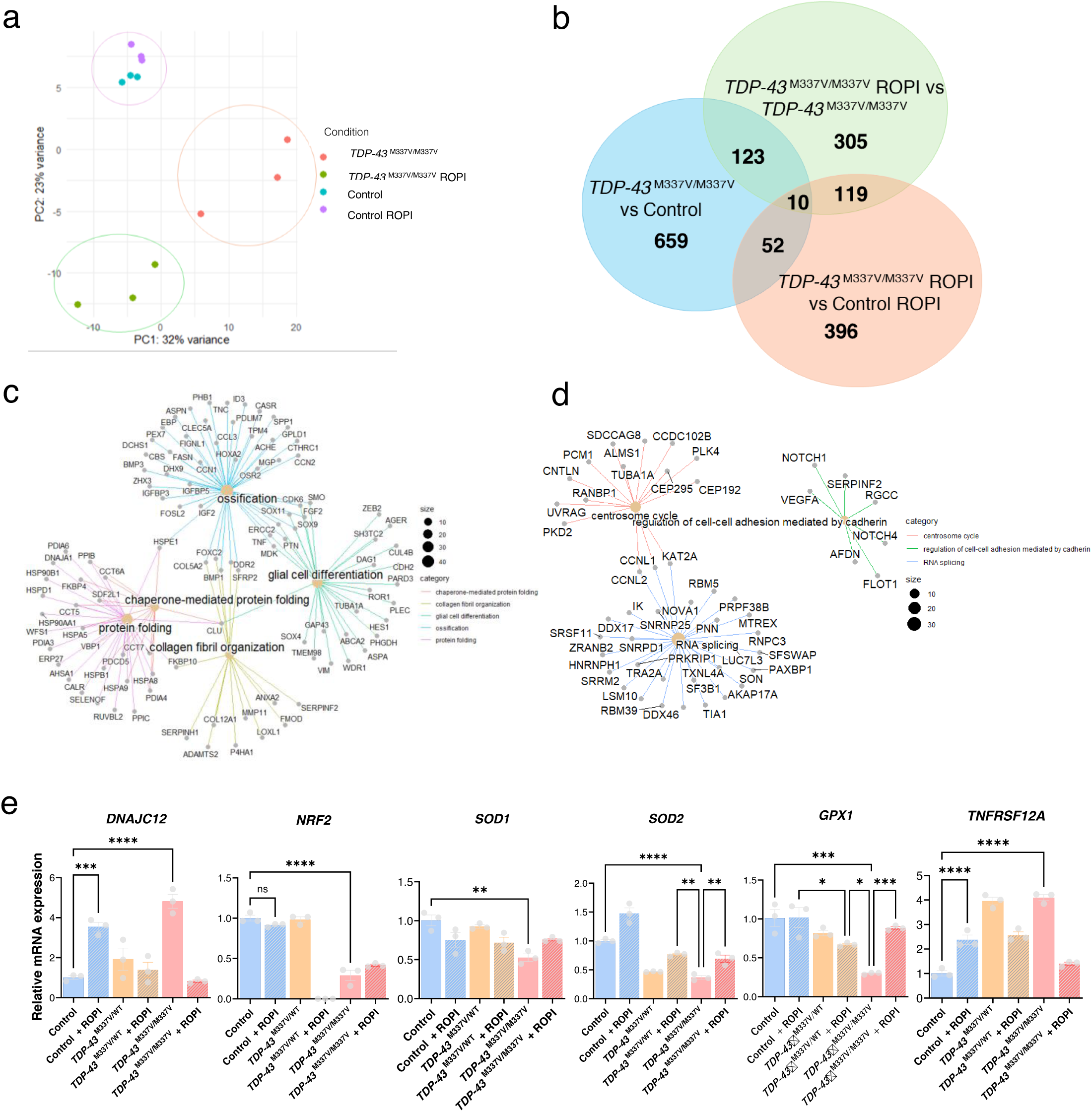
Global molecular profiling of TDP-43 iMGLs revealed transcriptional dysregulation in the TDP-43 mutant background and partial reversal of gene expression upon ropinirole treatment. (**A**) Principal component analysis results showing the transcriptomes of untreated and ROPI-treated control and *TDP-43*^M337V/M337V^ iMGLs. (**B**) Venn diagram showing the total number of DEGs between the two groups (adjusted p value < 0.05). (**C**) Bar graph showing the top significant gene ontology terms between *TDP-43*^M337V/M337V^ and the control. (**D**) Bar graph showing the top significant gene ontology terms between *TDP-43*^M337V/M337V^ ROPI and *TDP-43*^M337V/M337V^ untreated. (**E**) Validation of the top DEGs by qRT`‒`PCR and their reversal of expression upon ROPI treatment. The presented values are the means ± SEMs, n = 3 biological replicates.

Differential gene expression analysis (false discovery rate [FDR]<0.05) revealed 851 differentially expressed genes (DEGs) between untreated *TDP-43*^M337V/M337V^ iMGLs and control iMGLs (Fig. S2A, Table S2), establishing the transcriptional signature associated with the mutation. To assess the impact of ROPI, we compared ROPI-treated and untreated TDP-43^M337V/M337V^ iMGLs, which revealed 612 DEGs. Of these, 62 overlapped with both untreated mutant vs control comparison, and ROPI-treated mutant vs control comparisons (Fig. S2A, Table S2), while 396 were uniquely altered by ROPI treatment.

Importantly, 181 DEGs were shared between the ROPI-treated and untreated conditions, suggesting that ROPI partially counteracts mutation-associated transcriptional changes. Thus, the mutant vs control comparison defines the baseline perturbations caused by *TDP-43*^M337V/M337V^ mutation and the treated vs untreated comparison reveals how ROPI modifies this mutant-specific signature (Fig. 5b). Hierarchical clustering of the top 100 DEGs showed ROPI-treated *TDP-43*^M337V/M337V^ iMGLs clustered more closely with control iMGLs than with untreated mutants, suggesting that ROPI selectively modulates gene expression specifically in *TDP-43*^M337V/M337V^ iMGLs (Fig. S2B).

To further investigate the biological pathways affected by ROPI treatment, we performed gene ontology (GO) enrichment analysis. When we compared *TDP-43*^M337V/M337V^ iMGLs to control iMGLs, we observed significant enrichment in pathways related to protein folding, chaperone-mediated protein folding, extracellular matrix (ECM) organization, and glial cell differentiation (Fig. 5c, Supp. Table 3), suggesting that ROPI may modulate microglial structural integrity. Given the role of ECM in neuroinflammation, these findings support ROPI’s potential to enhance microglial function in ALS. Comparison of ROPI-treated mutant and control iMGLs revealed enrichment in RNA splicing and collagen metabolism pathways (Fig. 5d). Although TDP-43 regulates splicing, prior studies report conflicting reports for M337V-induced splicing defects [58,59]. Our data shows that ROPI alters splicing-related gene expression. We further examine *STMN2* transcripts and found reduced full-length and cryptic exon expression in *TDP-43*^M337V/M337V^ iMGLs. Cycloheximide treatment increased cryptic exon levels, indicating that these transcripts undergo nonsense-mediated decay (NMD) (Fig.S2C).

Given the role of oxidative stress in ALS, we examined whether ROPI modulates its related gene expression. RNA-seq revealed elevated oxidative stress-associated transcripts in untreated *TDP-43^M^*^337V/M337V^ iMGLs, which were normalized by ROPI comparable to the control levels (Fig. 5e), supporting ROPI’s antioxidant properties and neuroprotective effects. We also assessed cGAS, a cytosolic DNA sensor linked to neuroinflammation via the cGAS`‒`STING pathway. cGAS expression was upregulated in *TDP-43*^M337V/M337V^ iPSC-derived cells, consistent with previous reports of aberrant DNA sensing in ALS. Notably, treatment with ROPI restored cGAS expression to levels comparable to those of isogenic controls (Fig. S3A). Heatmap analysis of curated cGAS-STING genes showed broad upregulation trends in the *TDP-43^M337V/M337V^* iMGLs, which was attenuated by ROPI (Fig. S3B), suggesting ROPI may mitigate aberrant DNA sensing in ALS.

Taken together, our data show that ROPI treatment alters the transcriptomic profile of *TDP-43*^M337V/M337V^ iMGLs, partially restoring gene expression pattern toward control levels. ROPI also reduces oxidative stress, reinforcing its potential to mitigate inflammation-related pathology in ALS. Originally developed as a dopamine D2 receptor agonist for Parkinson’s disease, ROPI may have broader applications in neurodegenerative disease research and therapeutic development.

### Connectivity map analysis revealed dysregulation of the PI3K‒mTOR pathway in TDP-43 iMGL

To characterize the transcriptional landscape of TDP-43 iMGLs, we applied a computational framework that compares disease and control transcriptomes to identify compounds capable of reversing disease signatures. This strategy hypothesizes that reversing disease-associated transcriptional signatures may identify compounds mitigating TDP-43 pathology and offer therapeutic benefits in ALS. We generated an ALS-TDP-43 signature from significant DEGs and queried it against the CMAP database, focusing on THP-1 cell perturbagens due to their relevance to microglial biology. Top candidates included mammalian target of rapamycin (mTOR) inhibitors (PF-04691502, SB-2343, PF-05212384), PI3K inhibitors (idelalisib, taselisib, wortmannin, and GDC-0941), and the AKT inhibitor MK-2206 (Fig. S4A, Supp. Table 4). These results implicate the PI3K-Akt-mTOR signaling axis as a key regulator of the *TDP-43*^M337V/M337V^ iMGL transcriptome and suggest that its modulation may reverse disease-associated transcriptional dysregulation.

We further extended the analysis to include MNEU (iPSC-derived neuronal models) datasets in CMAP, where mTOR and PI3K inhibitors consistently ranked in the top 10 by mean connectivity score (Fig. S4B), reinforces the hypothesis that targeting the PI3K-Akt-mTOR pathway may be broadly applicable to various TDP-43-associated disease states.

We next assessed ROPI’s effects in TDP-43 iMGLs, generating ROPI-ALS-TDP-43 signature from DEGs and querying CMAP. The dopamine D4 receptor agonist ABT-724[60] showed a strong positive connectivity score with ROPI, aligning with its known mechanism and validating our connectivity-based approach. Notably, PI3K-Akt-mTOR inhibitors, including everolimus, GDC-0068, and SAR-245409, also showed strong positive connectivity with ROPI across THP-1 and neuronal datasets. These findings suggest that, in addition to dopaminergic modulation, ROPI may exert ancillary effects on the PI3K-Akt-mTOR signaling cascade (Fig. S4C, S4D). To further investigate this potential mechanism, we analyzed the expression patterns of genes related to the mTOR pathway. Heatmap visualization revealed that key mTOR-related transcripts were upregulated in TDP-43 iMGLs. Importantly, their expression levels were partially normalized by ROPI (Fig. S4E). The ROPI was also found within the CMAP signature list; however, its mean connectivity score was inferior to that of everolimus, an FDA-approved mTOR inhibitor currently used to treat various tumors[61,62] (Fig. S4F–I). Together, these findings uncover a previously unrecognized role for ROPI in modulating mTOR signaling in TDP-43-affected microglia, warranting further mechanistic investigation to elucidate the mechanism by which ROPI inhibits the mTOR pathway.

## Discussion

### *TDP-43* Mutation Induces Oxidative Stress and Autophagy Deficits in Microglia

TDP-43 is expressed at comparable levels across major brain cell types, but its role in microglia remains poorly understood. Using isogenic pairs of *TDP-43*^M337V/M337V^-mutant hiPSC-derived iMGLs, we found that the mutant iMGLs exhibit elevated oxidative stress, phagocytosis and cell-autonomous deficit in autophagy and mitophagy, potentially driving iron imbalance and altered ferritin levels. While autophagy dysregulation has been linked to TDP-43 pathology and ALS-related genes [63,64], its impact on microglia in a human context has been underexplored [58,65,66]. Our model isolates TDP-43 dysfunction and reveals that ROPI treatment partially reverses ALS-associated phenotypes [54–56]. This study offers the first evidence that ROPI may alleviate neuroinflammatory pathology in *TDP-43*^M337V/M337V^ microglia [58,65,67].

### Ropinirole Partially Restores Inflammatory and Transcriptomic Abnormalities

Oxidative stress resulting from aging or genetic mutations contributes to neurodegeneration [13,19,30,68]. We previously showed that ALS patient-derived motor neurons exhibit increased oxidative stress[54,55]. Here, we demonstrate that *TDP-43*^M337V/M337V^ iMGLs are similarly vulnerable, with oxidative stress evident from early development, potentially compounding disease risk. While oxidative damage typically targets neurons, our findings highlight microglial susceptibility.

Microglia respond to cellular insults via phagocytosis. Contrary to reports of impaired microglial phagocytosis in ALS models[31,32,34], TDP-43 iMGLs showed increased phagocytic activity suggesting compensatory or stage-dependent response. This aligns with early-stage microglial phenotypes, and prior evidence of phagocytic marker upregulation in ALS brains [29,30] Similar hyperphagocytic states have been reported in other neurodegenerative conditions, including Alzheimer’s disease, Parkinson’s disease[56] and frontotemporal dementia[57], where excessive phagocytosis correlates with neuronal or synaptic loss linked to subtype-specific vulnerabilities [58]. While phagocytosis support homeostasis, excessive or a misdirected activity, especially via a faulty receptor signaling, can harm viable neurons. Our data suggest that TDP-43 iMGLs may intrinsically adopt a hyperphagocytic state, underscoring the need to balance microglial activity in therapeutic strategies.

*TDP-43* iMGLs showed impaired autophagic flux, and reduced mitophagy, mirroring deficits seen in *C9orf72 i*PSC-derived iMGLs and leading to poor clearance of protein aggregates and damaged mitochondria [31]. Similar mitochondrial abnormalities have been reported in TDP-43 overexpression models [70–72]. Autophagy dysregulation is a known contributor to ALS, and inducers can mitigate TDP-43 pathology in mouse models. Since autophagy is suppressed by mTOR and TDP-43 helps stabilize ATG7 mRNA, its mutation disrupts oxidative stress response, autophagy and mitophagy, driving neuroinflammation. Notably, ROPI treatment failed to restore these deficits in *TDP-_43_*M337V/M337V _iMGLs._

Altered cytokine profiles in ALS patients underscore the role of neuroinflammation [73]. In *TDP-43*^M337V/M337V^ iMGLs, we observed elevated IL-8 and reduced IL-6 and IL-10, indicating a shift toward proinflammatory state and impaired resolution of inflammation. While C9orf72 and VCP iMGLs showed increased IL-6, FUS models show minimal cytokine changes [31,34,35]. We also found modest transcriptional upregulation of the cGAS-STING pathway, consistent with its emerging role in DNA damage-induced immune activation and neurodegeneration [74–79]. Aberrant activation of this pathway may reflect a maladaptive response to DNA damage or oxidative stress, contributing to disease progression. In addition, ROPI treatment modulated the expression of several upregulated transcripts within this pathway, warranting further investigation to clarify the mechanistic relevance and therapeutic potential of cGAS-STING signaling in ALS.

Ferritin light chain (FTL) levels were significantly reduced in *TDP-43*^M337V/M337V^ iMGLs, suggesting disrupted iron metabolism that may worsen oxidative stress and inflammation [29,30]. While ROPI restored ferritin heavy chain (FTH1) transcript levels, it did not normalize FTL. Given their distinct roles in iron handling, further study is warranted to clarify how TDP-43 mutations impair iron homeostasis. Ferritin protects against ROS by sequestering iron, and its dysregulation may increase vulnerability to oxidative damage. Reduced *FTH1* in TDP-43 knockdown neuron [73] supports a model in which iron imbalance drives neurodegeneration via ROS and mitochondrial dysfunction.

### mTOR Pathway Modulation and Therapeutic Implications

RNA-seq analysis revealed that the M337V mutation in TDP-43 significantly altered the transcriptomic landscape of iMGLs, with enrichment in pathways related to protein folding, ECM organization and glial differentiation. ROPI treatment partially reversed these changes, particularly affecting ECM and collagen metabolism. Dysregulation of heat shock proteins (HSPs), such as *HSPA5, HSPA8*, and *HSPA9* suggests impaired protein quality control, a key feature in ALS pathology. Collagen-related changes and MMP activity point to ROPI’s role in modulating ECM structure and inflammation.

Additionally, ROPI influenced RNA splicing-related genes, relevant given TDP-43’s role in repressing cryptic exons [80,81]. Several studies have reported that TDP-43 Q331 or M337V has prominent splicing defects, as shown by cryptic exon expression of *STMN2*. However, the M337V mutation has been demonstrated only in an overexpression mouse model that exhibits a splicing defect in the spinal cord[82]. We observed reduced full-length and cryptic *STMN2* transcripts in *TDP-43*^M337V/M337V^ iMGLs, likely due to NMD. Cycloheximide treatment restored cryptic *STMN2* expression, confirming that the M337V mutation disrupts splicing via TDP-43 loss of function. Together, these data support that *STMN2* splicing deregulation is due to the loss of TDP-43 function caused by the M337V mutation.

In FUS P525L iPSC-derived motor neurons, PI3K/mTOR inhibition activates autophagy and mitigates stress granule pathology, conferring neuroprotective effects[83]. Similar PI3K/Akt inhibition improves survival in ALS mouse models[84,85], and rapamycin, an mTOR inhibitor alleviates autophagic dysfunction and ALS-associated phenotypes in a TDP-43 mouse model [86]. Consistent with these findings, our drug repurposing analysis identified PI3K-Akt-mTOR inhibitors as top candidates for reversing the transcriptomic signature of TDP-43 iMGLs, in both THP-1 cells and iPSC-derived neuronal signatures [87]. CMAp analysis of ROPI-treated iMGLs revealed hits along PI3K-Akt-mTOR inhibition axis, suggesting ROPI may modulate this pathway, This is further supported by the partial reversal of mTOR-related gene expression following ROPI treatment. ROPI treatment has been shown to enhance p70S6K expression, a downstream effector of the mTOR pathway, indicating mTOR activation through dopamine receptor signaling[88].

Together, these findings uncover a previously underappreciated role for microglia in ALS pathogenesis, driven by cell-autonomous TDP-43 dysfunction. The convergence of oxidative stress, autophagy and mitophagy deficits, cytokine imbalance and iron dysregulation underscores the multifaceted contribution of microglia to neurodegeneration. Importantly, ROPI not only exerts neuroprotective effects in motor neurons but also modulates key inflammatory and metabolic pathways in microglia, including PI3K-Akt-mTOR signaling. Future investigation of TDP-43 iMGL and its interactions with motor neuron or organoid coculture and xenotransplantation models may help to further elucidate synaptic pruning and microglial phenotypes that are relevant to neurodegeneration. Overall, the present study identifies downstream cellular effects regulated by TDP-43 mutation specifically in microglia. By illuminating early-stage microglial vulnerabilities and identifying actionable therapeutic targets, this study advances our understanding of ALS and highlights the potential of targeting neuroinflammation and microglial dysfunction to reshape disease trajectory.

### Limitation

The current study utilizes a single isogenic iPSC line, with the homozygous mutant serving as the primary model for our experimental assays. While this approach provides a controlled system to probe mutation-specific effects, it does not capture the heterogeneity present in patient-derived or heterozygous backgrounds. Nonetheless, we used this model to study the functional impairment associated with biallelic TDP-43 mutation and its impact on microglial dysfunction, providing a controlled system to probe mutation-specific effects. We acknowledge this as a limitation and recognize that incorporating additional iPSC lines, including heterozygous mutations and patient-derived samples, will be important to strengthen the translational relevance and generalizability of our conclusions.

### Conclusion

In summary, this study establishes a human isogenic iPSC-derived microglial model of TDP-43 pathology in ALS and identifies ROPI as a candidate therapeutic compound that mitigates oxidative stress, inflammatory dysregulation, and transcriptomic abnormalities. These findings broaden the cellular scope of ROPI from motor neurons to microglia, emphasizing the need to target nonneuronal mechanisms in ALS. Our results further support PI3K-Akt-mTOR signaling as a therapeutic axis and underscore the value of iMGL-based platforms for ALS drug discovery.

## Declarations

### Ethics approval and consent to participate

Not applicable

### Consent for publication

### Availability of data and materials

The raw sequencing data used in this study are available at GEO with accession numbers. GSE288292.

### Competing interests

H.O. is the cofounder and an SAB of K Pharma, Inc. The other authors declare that they have no known competing financial interests or personal relationships that could have appeared to influence the work reported in this paper.

### Funding

This study was supported by grant support from the Japan Society for the Promotion of Science (JSPS) (KAKENHI Grant No. JP21H05278, JP22K15736 and JP25H00007 to S.M.), the Japan Agency for Medical Research and Development (AMED) (Grant No. JP23bm1123046, JP23kk0305024, JP25ek0109811 to S.M., JP21wm0425009, JP22bm0804003, JP22ek0109616, JP23bm1423002, JP25wm0625519 to H.O.), the Daiichi Sankyo Foundation of Life Science, the UBE Academic Foundation, the Kato Memorial Trust for Nambyo Research, Japan Intractable Diseases (Nanbyo) Research Foundation (2024A04), the Inamori Foundation and Kanagawa Institute of Industrial Science and Technology (KISTEC) to S.M. during the conduct of the study.

### Authors’ contributions

Conceptualization was performed by H.O., S.M., S.T. and K.H.U. Experiments were performed by K.H.U. and T.K. Analysis was performed by S.M., N.T., H.O., and K.H.U. Resources were provided by H.O., H.W., K.O., S.M., and Y.M. Writing of the original draft was performed by S.M., H.O., and K.H.U. Review and editing were performed by H.O., S.M., T.K., Y.M., H.W., S.T., K.O., and K.H.U. Funding acquisition was performed by S.M., Y.M., and H.O. Supervision was performed by S.M., H.W., S.T., Y.M., and H.O.

## List of abbreviations

ALS: amyotrophic lateral sclerosis
CMAP: Connectivity map
DEG: differentially expressed genes
ECM: extracellular matrix
GO: Gene ontology
GSEA: Gene set enrichment analysis
HSP: heat shock protein
hiPSC: human induced pluripotent stem cells
iMGL: induced microglial-like cells
mTOR: mammalian target of rapamycin
NMD: nonsense-mediated decay
ROPI: Ropinirole
TDP-43: TAR DNA binding protein 43

## Acknowledgments

We thank the Okano laboratory members for their invaluable suggestions and input. We thank the Collaborative Research Resources at Keio University School of Medicine for their technical assistance.

## Additional Files

Supplementary Figures 1--3.

Supplementary Tables 1--4.

