## Supplemental Figures for "Ropinirole hydrochloride mitigates oxidative stress and neuroinflammation via the PI3K–mTOR pathway in TDP-43 hiPSC-derived microglial-like cells"

Supplementary Figure 1

#### Figure S1A.

(Left) Control and TDP-43$^{M337V/M337V}$ express myeloid-lineage marker (IBA1) and microglial specific marker (P2RY12) upon maturation (Scale bar: 50 μm). (Right) Quantification of IBA1+ cells over total number of DAPI-stained nuclei from two independent differentiations and quantified from a total of 12,000 nuclei

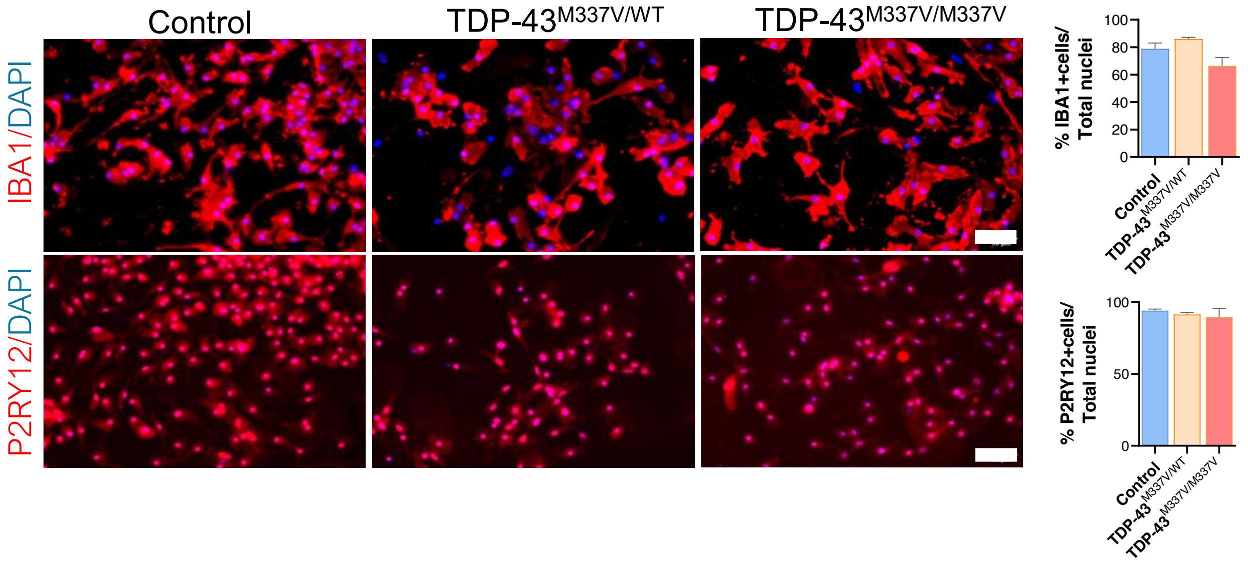


#### Figure S1B.

Time-lapse imaging for incucyte analysis for pHRodo-Zymosan beads phagocytosis assay in Control and TDP-43$^{M337V/M337V}$ iMGLs (correspond to Fig.2B)

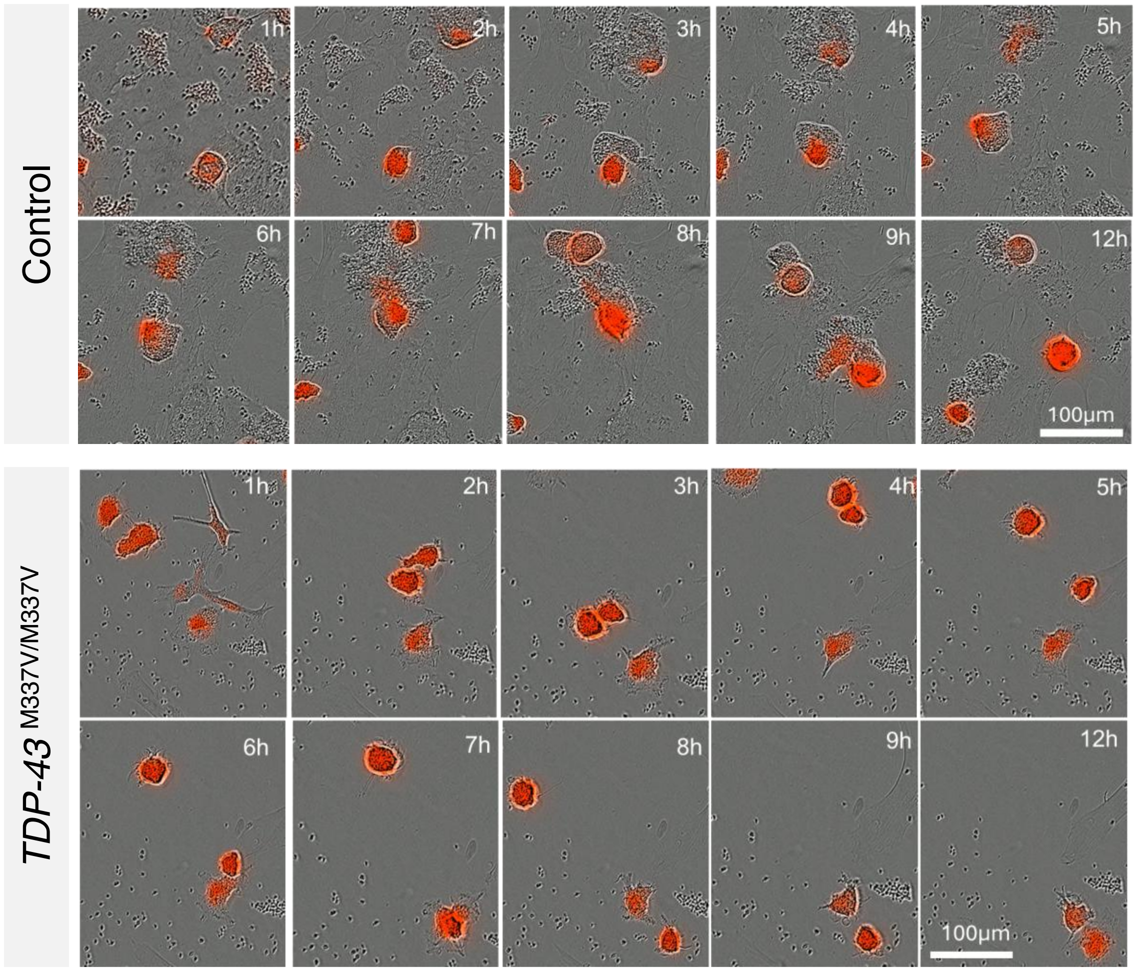

### Supplementary Figure 2

#### Figure S2A.

Volcano plot showing significant DEGs between TDP-43$^{M337V//M337Vvs}$ Control (left) and ROPI-treated TDP-43$^{M337V//M337Vvs}$ Control (right).

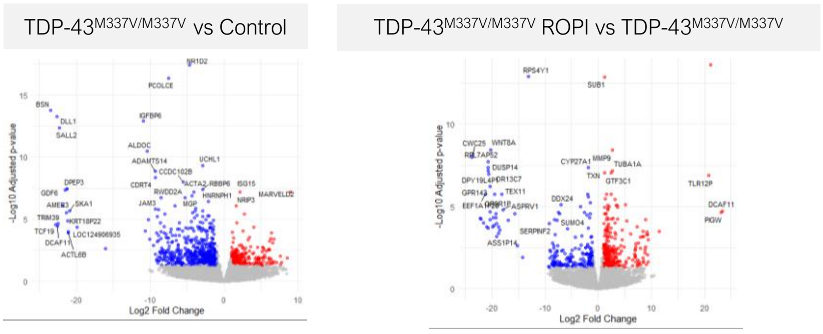


#### Figure S2B.

Heatmap representation and hierarchical clustering of all the samples. Rows are clustered using Eucledian method.

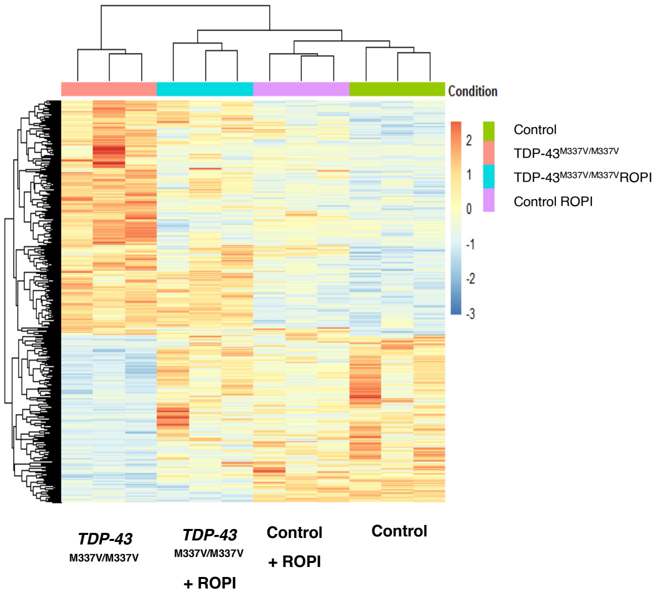


#### Figure S2C.

Full length and cryptic exon *STMN2* expression are significantly reduced in TDP-43$^{M337V/M337V}$ iMGL, but upon translation inhibition with cycloheximide its expression is significantly increased suggesting NMD mechanism

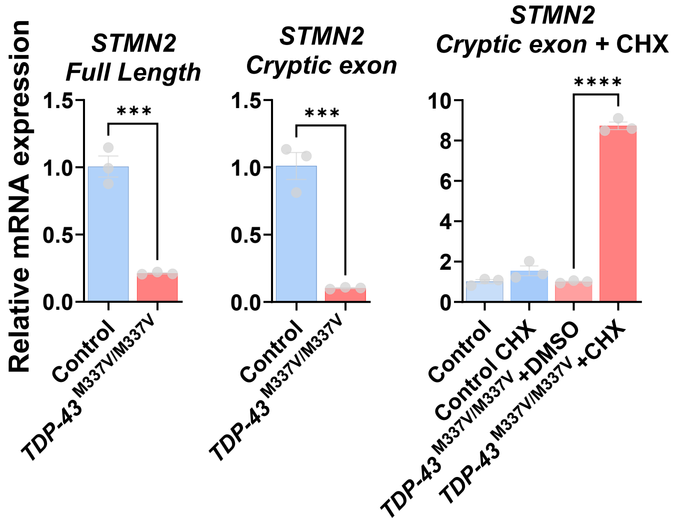

#### Supplementary Figure 3

##### Figure S3A.

Counts per million values of cGAS expression derived from RNAseq analysis of isogenic TDP-43$^{M337V/M337V}$ iMGLs following ROPI treatment.

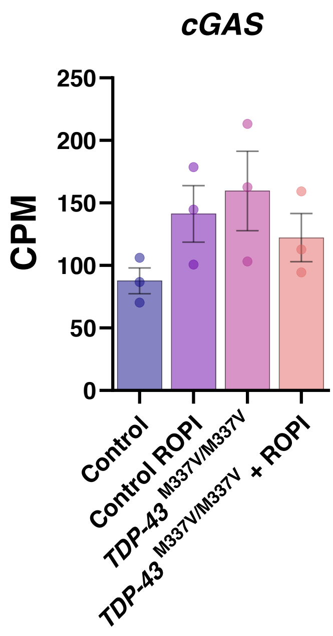


##### Figure S3B.

Heatmap showing differential expression of cGAS-STING pathway-transcripts in isogenic TDP-43$^{M337V/M337V}$ iMGLs following ROPI treatment. Data were normalized and log-transformed.

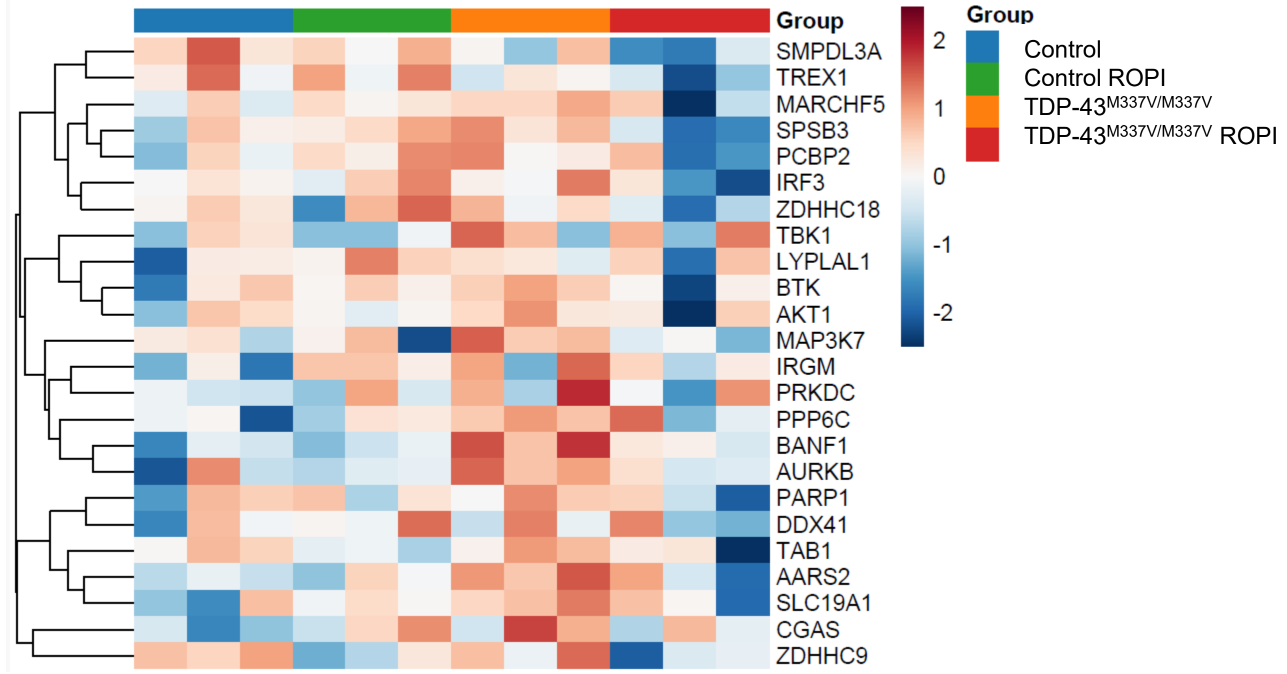

### Supplementary Figure 4

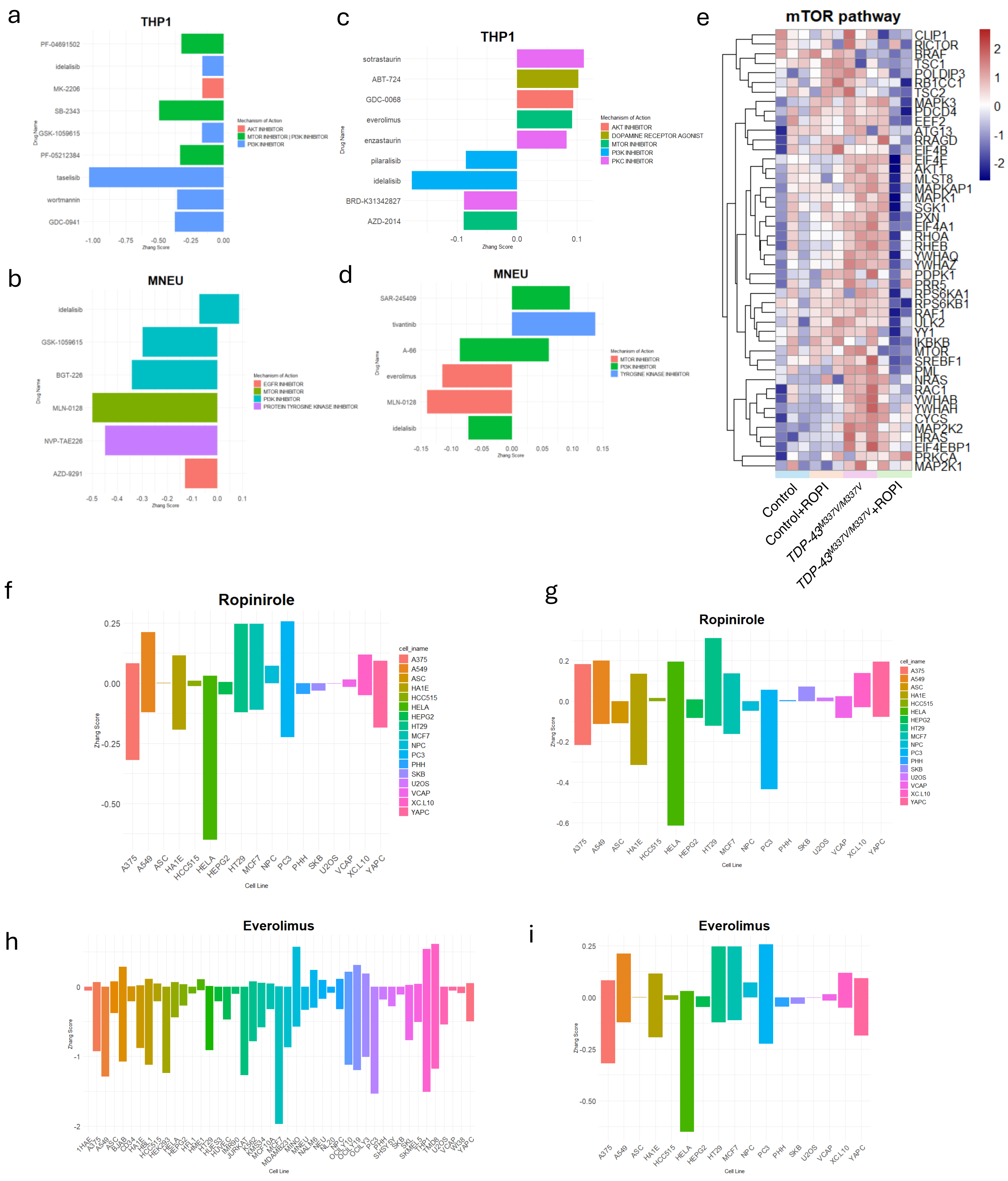


**Figure S4. Connectivity Map (CMap) analysis identified PI3K-Akt-mTOR inhibition reverses the transcriptional signature of TDP-43 iMGL.** (**A**) Top rank of transcriptional reversal from THP1 signature in CMap for TDP-43 M337V iMGL vs Control. (**B**) Top rank of transcriptional reversal from iPSC-derived neuronal signature in CMap TDP-43 M337V iMGL vs Control. (**C**) Top rank of transcriptional reversal from THP1 signature in CMap for Ropi-treated TDP-43 M337V iMGL vs untreated TDP-43 iMGL. (**D**) Top rank of transcriptional reversal from iPSC-derived neuronal signature in CMap for Ropi-treated TDP-43 M337V iMGL vs untreated TDP-43 iMGL. (**E**) Heatmap visualization of mTOR pathway component generated by z-score normalization. (**F-G**) Ropinirole CMap signature across different cell lines for TDP-43 M337V iMGL vs Control (**F**), and Ropi-treated TDP-43 M337V iMGL vs untreated TDP-43 iMGL (**G**). (**H-I**) Everolimus CMap signature across different cell lines for TDP-43 M337V iMGL vs Control (**H**), and Ropi-treated TDP-43 M337V iMGL vs untreated TDP-43 iMGL (**I**)
